# A scalable long-term *in vitro* model of human colonic epithelium with continuous barrier function

**DOI:** 10.1101/2025.09.24.677447

**Authors:** Ashley J. Cuttitta, Sophia R. Meyer, Tristan Frum, Bhargavasai Gunuguntla, Angeline Wu, Sha Huang, Jack Morgan, Michael K. Dame, Jonathan Z. Sexton, Jason R. Spence

**Affiliations:** Department of Internal Medicine, The Division of Gastroenterology and Hepatology, The University of Michigan Medical School, Ann Arbor, MI 48109, USA; Translational Tissue Modeling Laboratory Core Facility, The University of Michigan Medical School, Ann Arbor, MI 48109, USA; Department of Medicinal Chemistry, College of Pharmacy, University of Michigan, Ann Arbor, MI 48109, USA; Department of Cell and Developmental Biology, The University of Michigan Medical School, Ann Arbor, MI 48109, USA; Department of Biomedical Engineering, The University of Michigan Medical School, Ann Arbor, MI 48109, USA

**Keywords:** 2D, high-content imaging, organoid, TEER

## Abstract

**Background & Aims:** Human *in vitro* models of differentiated intestinal epithelium are valuable research tools for discovery and translational medicine. Primary adult colonic epithelium barrier function is difficult to assess in a 2-dimensional (2D) *in vitro* format for more than a few days due to rapid cellular turnover. We developed a transwell model of the colonic epithelium that maintains proliferative and differentiated cell types, allowing long-term culture.

**Methods:** Using 3-dimensional (3D) primary human colonoids, we seeded dissociated epithelium onto 96-well transwell membranes and added different media that was growth factor specific to apical or basolateral transwell chambers. Barrier integrity was assessed using a variety of stimuli and TEER was continuously monitored in real time for up to 2 weeks, followed by quantitative single-cell image analysis using high-content imaging and deep-learning-based cell segmentation.

**Results:** Epithelium cultured up to 30 days sustained a time-dependent differentiation, leading to the establishment of a monolayer with proliferative stem/progenitor and differentiated cell types. While monitoring continuous TEER across multiple donor organoids, barrier challenge assays recorded donor-specific responses to IFN-γ, TNF-α, *Clostridium difficile* toxin B (TcdB), and ethylenediaminetetraacetic acid (EDTA). Endpoint immunofluorescence analysis of tight junctions and whole cell staining identified unique cellular features among treatment groups which significantly correlated with TEER, including cell-cell orientation (e.g., adjacent cell mean angle).

**Conclusions:** We demonstrate a high-throughput *in vitro* model of the homeostatic human colonic epithelium that maintains a long-term functional barrier. This model can be used to investigate complex epithelial barrier responses over time.

## Introduction

*In vitro* models, such as 3-dimensional (3D) organoids or 2-dimensional (2D) monolayers of human cells are excellent ways to generate translational data from human tissue, and are an alternative to animal models, which have historically low success rates in the development of effective therapies during the preclinical stage [1, 2]. Well characterized *in vitro* models of human intestinal epithelium derived from expandable human donor intestinal stem cells (ISCs) [3] recapitulate many aspects of the form and function of the *in vivo* intestinal epithelium, and are valuable resources to study biological processes, disease mechanisms, and develop therapeutics.

The luminal surface of the intestine is comprised of the epithelium, a selectively permeable barrier of polarized cells organized into crypts and villi. The epithelium provides protection to intestinal sublayers from luminal contents while having both absorptive and secretory functions [4, 5]. The intestinal epithelium is also highly regenerative, containing ISCs at the base of the crypts that proliferate, migrate and terminally differentiate, replenishing the epithelial surface every 3-5 days with the capacity to repair damage induced by barrier disruption. The expansion of human colon ISCs as 3D organoids [3, 6] or 2D monolayers [7, 8] has enabled the development of a broad spectrum of *in vitro* models containing *in vivo*-like epithelium. Complex models of human colon epithelium like organ-on-a-chip mini-colons [9] and collagen scaffold crypt arrays [10, 11] integrate elements of the epithelial environment by recreating the 3D architecture of the crypt with representative cellular diversity including goblet cells that secrete mucin to form a mucus layer. However, these models are not easy to reproduce without bioengineering expertise and resources. Transwell inserts made for typical cell culture dishes are a commercially available, simple and widely used way to model epithelium with the benefit of open access to the epithelium’s apical side. ISCs are seeded and differentiated on the 2D porous transwell membrane coated with extracellular matrix protein. The semipermeable membrane with attached cells hangs from the bottom of a plastic cup into a receiver well, creating apical and basolateral compartments. These compartments allow easy manipulation of culture medium and additives exclusively to either side of the polarized epithelium, supporting versatile experimental design like co-culture [12–15] and drug screening [16] that can be scaled up for high-throughput assays. In addition, most transwell inserts are designed for use with an epithelial volt/ohm meter (EVOM), having a small opening at the top of the transwell cup for the insertion of an electrode into the apical and basolateral compartments. A non-invasive electrical current is passed over the epithelium allowing for the measurement of transepithelial electrical resistance (TEER). TEER is proportional to the adherence of epithelial cells to each other by tight junctions that form the epithelial barrier and control the passage of ions across it [17, 18]. Generally, the formation of functional TEER requires a fully differentiated epithelium, which corresponds to a loss of proliferative stem/progenitor cells as they differentiate [19, 20]. Thus, *in vitro* barrier function and TEER is often short-lived, making the long-term study of a physiological epithelial barrier function difficult.

A primary goal of the current work was to overcome the limitation of short-term TEER, and to determine if we could maintain a homeostatic epithelium that possessed both proliferative progenitors and differentiated epithelium in long-term 2D culture. To do this, we modified our well established working protocol for terminally differentiated, short-term colon epithelial monolayers on 24 well transwells [21–23], which has been adapted from previously published methods [12, 24–30]. We substituted the medium that is usually used to induce differentiation, and instead used ISC supporting medium in the basolateral (lower) chamber of the transwell system and growth factor replete medium in the apical (upper) chamber to encourage cellular differentiation. This combination supported a polarized long-term epithelial monolayer with columnar shaped cells that maintained proliferating progenitor cells and differentiated cell types with a functional barrier for 3-4 weeks, the typical time that we stopped the culture experiments. Next, we created long-term epithelial cultures in 96 well transwells for use with a high-throughput continuous TEER monitoring system [31] and examined epithelial responses to barrier challenge by regular addition of pro-inflammatory cytokines interferon gamma (IFN-γ), tumor necrosis factor alpha (TNF-α), both combined, *Clostridium difficile* toxin B (TcdB) or ethylenediaminetetraacetic acid (EDTA) for >1 week. Finally, at the end of the TEER experiment, we fixed, stained and analyzed the monolayer post treatment using immunofluorescence (IF), followed by quantitative single-cell resolution image analysis using high-content imaging and deep-learning-based cell segmentation. Analysis of tight junction proteins and whole cell staining combined with quantitative analysis identified unique cellular features among treatment groups which significantly correlated with TEER, allowing us to identify a cellular phenotype that correlates with a strong epithelial barrier. Together, this work transforms the primary human colonic epithelium into a robust, high-throughput discovery engine for evaluating barrier-modulating drugs and biologics, and enables mechanistic dissection of barrier dysfunction in human disease over time and at scale.

## Results

### Long-Term (LT) epithelium forms in vivo-like cell morphologies and functional barrier with proliferating and differentiated cells for up to 30 days

We sought to design a human colonoid derived long-term (LT) epithelium on a transwell that maintains a functional barrier. Previously, a homeostatic epithelium was achieved using mouse or human colonoids with stem cell supporting medium (L-WRN conditioned medium [32]) in the transwell basolateral compartment and a medium-free apical compartment, creating an air-liquid interface [33, 34]. In this model, submersion of the cells by adding media to the upper chamber caused an injury response [33]. Others have shown that a basolateral ISC supporting medium with basal medium in the upper chamber could be used to culture mouse small intestine derived 2D epithelial monolayers, producing a steadily increasing TEER for 20 days [35]. Therefore, we used ISC expansion medium (see methods) in the bottom chamber and growth factor replete medium (2D differentiation medium (2DM)) in the upper chamber, allowing us to maintain an LT epithelium for up to a month. This also permitted the use of an electrode to measure TEER at any time. Histological evaluation of transwell cultures revealed that the long-term (LT) epithelium model develops a taller, more *in vivo*-like columnar epithelium along with goblet-like cell morphologies compared to terminally differentiated short-term (ST) epithelial monolayers cultured completely in 2DM for 7 days (**Figure 1A**). We quantitated columnar cell heights from cross sections from the ST and LT monolayers at selected time points and the *in vivo* colon epithelium at the crypt at base, middle and top levels. The LT monolayers have a significantly increased mean cell height compared to the ST epithelium, while the mean cell height of the LT epithelium at day 30 was similar to the mean cell height measured at the top of the adult crypt in an *in vivo* tissue section (**Figure 1B**). This finding shows that the LT epithelium cell heights at day 30 are most like the differentiated and luminal region of the in *vivo* crypt (**Figure 1B**).

**Figure 1.**
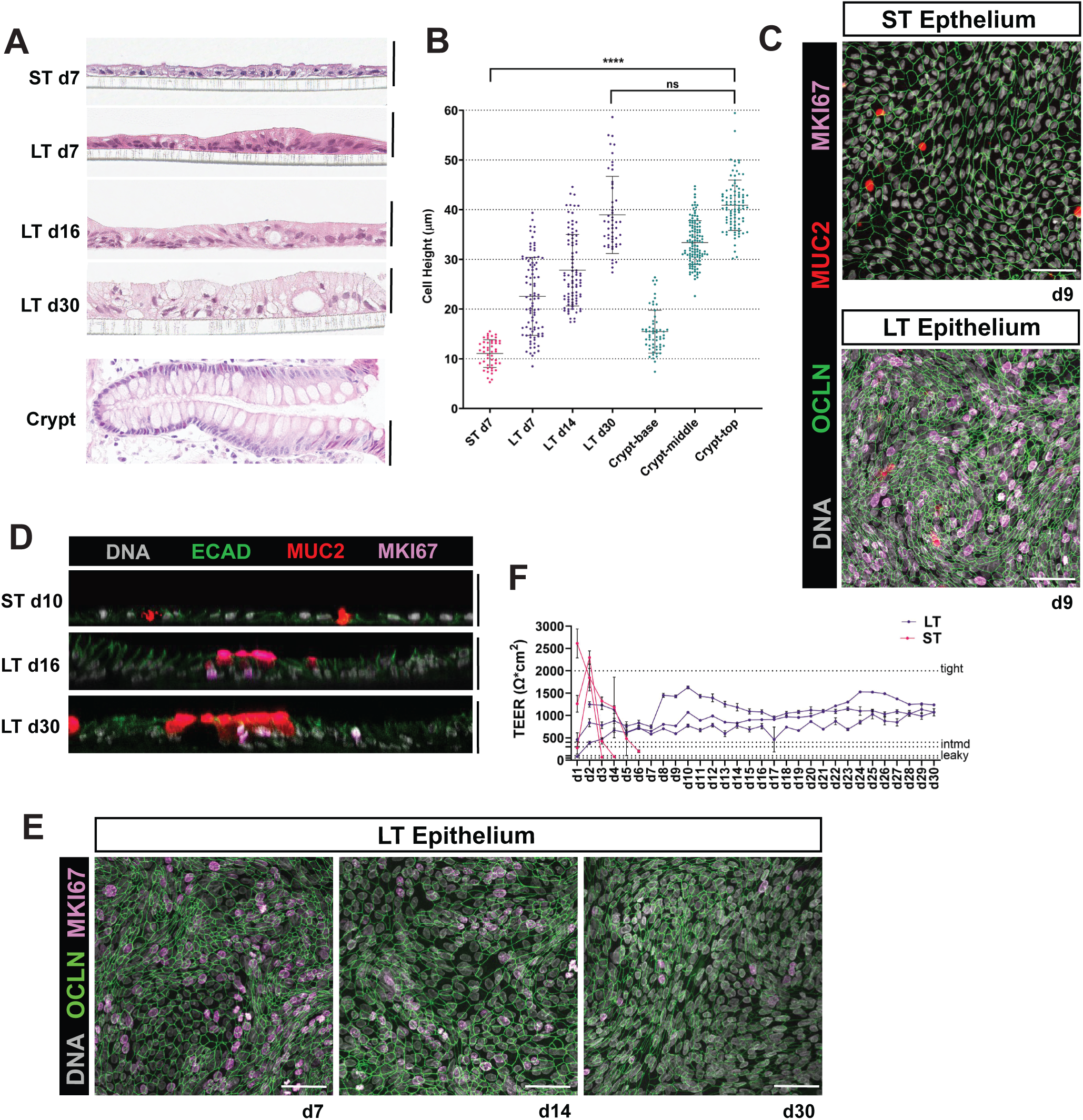
*In vitro* primary human colonoid derived LT epthelium characterization reveals *in vivo* like cell morphology, proliferative and differentiated cell populations, and functional barrier up to 30 days. **A.** Representative images of histological cross-sections from *in vitro* ST epithelium fixed on day 7 in culture, LT epithelium fixed on days 7, 16 and 30 in culture, and adult *in vivo* colon crypt stained with Hematoxylin and Eosin (H&E). Images were taken at 40X with an Leica Aperio slide scanner. Scale bars represent 50 mm. **B.** Quantitation of the *in vitro* ST, LT, and *in vivo* crypt epithelial cell heights. Data shown as mean ± SD of three technical replicates (n ≥ 3) with total number of cells measured per group: ST epithelium day 7 (n = 47), LT epithelium day 7 (n = 100), LT epithleium day 14 (n = 88), LT epithe-lium day 30 (n = 33), crypt base (n = 59), crypt middle (n = 118), crypt top (n = 92). Statistical signifiance was calculated using Welch’s *t*-test; *p* < 0.0001 (****) or *p* = 0.1135 (ns). **C.** Representative maximum intensity projection images of whole-mount IF stained ST epithelium (top) and LT epithelium (bottom) at day 9. Z-stack images were taken at 40X with a Nikon A1 light scanning confocal microscope. Scale bars represent 50 mm. **D.** Representative orthoganol view images of whole-mount IF stained ST epithelium at day 10 (top), LT epithelium at day 16 (middle), and LT epithelium at day 30 (bottom). Z-stack images were taken at 40X with a Nikon A1 light scanning confocal microscope. All scale bars represent 40 mm. **E.** Representative maximum intensity projection images of whole-mount IF stained LT epithelium at day 7 (left), day 14 (middle), and day 30 (right). Z-stack images were taken at 40X with a Nikon A1 light scanning confocal microscope. Scale bars represent 50 mm. **F.** Daily TEER measurement graphed as mean ± SD from six independent experiments (n ≥ 3 transwells per experiment) using Colon 88 ST epithelium (pink) or LT epithelium (purple). TEER classifications indicating tight (2000 Ω*cm²), intermediate (intmd; 300-400 Ω*cm²), and leaky (50-100 Ω*cm²) epithelial barrier are shown by dotted lines corresponding to respective Ω*cm².

We observed an epithelial barrier protein, OCCLUDIN (OCLN) by IF staining in both ST and LT epithelial cultures at day 9, whereas the proliferation marker MKI67 was detected only in the LT epithelial culture (**Figure 1C**) at the same timepoint. An orthogonal view of the ST epithelium shows short, E-CADHERIN (ECAD) bordered cells with MKI67 negative nuclei while LT epithelium at later time points (day 16 and day 30) show columnar ECAD stained cells with MKI67 positive nuclei in a pseudostratified pattern with proximity to MUCIN2 (MUC2) positive goblet cells at the luminal surface (**Figure 1D**). LT epithelial culture maintained an intact, OCLN positive barrier with MKI67 positive nuclei up to 30 days (**Figure 1E**). We compared ST and LT epithelial barrier function by collecting manual TEER measurements daily. Referring to gastrointestinal epithelia TEER classifications [36–38], the ST epithelium quickly develops a temporary tight (2000 Ω*cm^2^) with intermediate (300-400 Ω*cm^2^) and leaky (50-100 Ω*cm^2^) barrier measurements only a few days later, reflecting the ability of these cells to rapidly differentiate and form a barrier, followed by cell turnover and reduced barrier function. LT epithelium quickly develops a barrier that is classified between intermediate and tight, persisting for up to 30 days of measurement and stable among n = 3 different biological replicates (**Figure 1F**).

### Time-series single nucleus RNA sequencing of LT epithelium cultures reveals maintenance of TA-like cells and maturation of colonic cell lineages over time

In order to interrogate the cell populations present in the LT monolayer across 30 days *in vitro*, the epithelium was collected from transwell membranes on day 7 (early), day 14 (mid) and day 28 (late), cryopreserved, and then nuclei were isolated for single nucleus RNA sequencing (snRNA-seq) to determine the cell types that were present in LT culture across time. We used a label transfer approach to annotate LT monolayer cell types using a publicly available snRNA-seq dataset of *in vivo* ascending colon tissue served as the reference [39, 40]. *In vivo* reference data possessed a defined cellular differentiation trajectory from stem cell > transient amplifying 2 (TA 2) > transient amplifying 1 (TA 1) > immature colonocytes > colonocytes [40]. The early monolayer possessed cells expressing stem cell markers, though it also contained large proportions of cycling TA 2, cycling TA 1, and immature goblet cells. Mid and late epithelium shifted to predominately cycling TA 2 and goblet cells, indicative of goblet cell maturation over time (**Figure 2A**). Small populations of BEST4^+^ cells, non-cycling TA 1 cells, immature colonocytes, and colonocytes were measured at early, mid and late timepoints (**Figure 2A**). We did not identify any non-cycling TA 2, Tuft, or Enteroendocrine cells *in vitro* (**Figure 2A, 2B**). In the LT model, stem cells expressed canonical markers *SMOC2*, *RGMB*, and *ASCL2* (**Figure 2C**) [40–44]. A high percentage of immature goblet cells express *TFF3, MUC5B* and the crypt base marker *IGFBP2* [45], while mature goblet cells express high *MUC4,* and low *IGFBP2* and *MUC5B*. Both Mucin genes *MUC4* and *MUC5B* have previously been shown to be enriched in colon tissue [42] (**Figure 2C**). *KRT8* and *KRT18* expression in immature goblet and BEST4^+^ colonocytes may suggest active cell differentiation [46, 47] (**Figure 2C**). *MUC2*, a robust marker of goblet cells that encodes the primary secreted mucin (MUC2) from goblet cells in the colon, was not detected by snRNA-seq in LT epithelium goblet cell populations (**Figure 2C**); however, we find abundant MUC2 positive cells by whole-mount IF staining (**Figure 1C, 1D**), suggesting that mRNA levels of MUC2 are not detected in our snRNA-seq data. We validated MUC2 expression by immunohistochemical (IHC) labeling of LT epithelium sections alongside colon crypt sections. We also found that LT monolayer epithelium increased MUC2 expression with addition of Vasoactive Intestinal Peptide (VIP) to the basolateral medium as previously described (**Supplementary Figure 2**) [48]. Based on the discrepancy between mRNA and protein staining, it is likely that the failure to detect *MUC2* in snRNA-seq may be a limitation of this sequencing technology. Genes expressed in the differentiated colonic epithelium including *ATP8A1* [49], *KRT20* [50], and *CECAM1* [40] are similarly expressed *in vivo* and *in vitro* colonocyte populations (**Figure 2C**). Similarly, genes coding for ion transport in luminal epithelial cells, including *SLC26A3* [51], *CLCA4* [52], and *NR1H4* [53], along with tight junction genes *TJP1*, *OCLN*, and *CLDN4*, important in the maintenance of colonic epithelial barrier [54, 55], were also similarly expressed *in vivo* and *in vitro* (**Figure 2C**).

**Figure 2.**
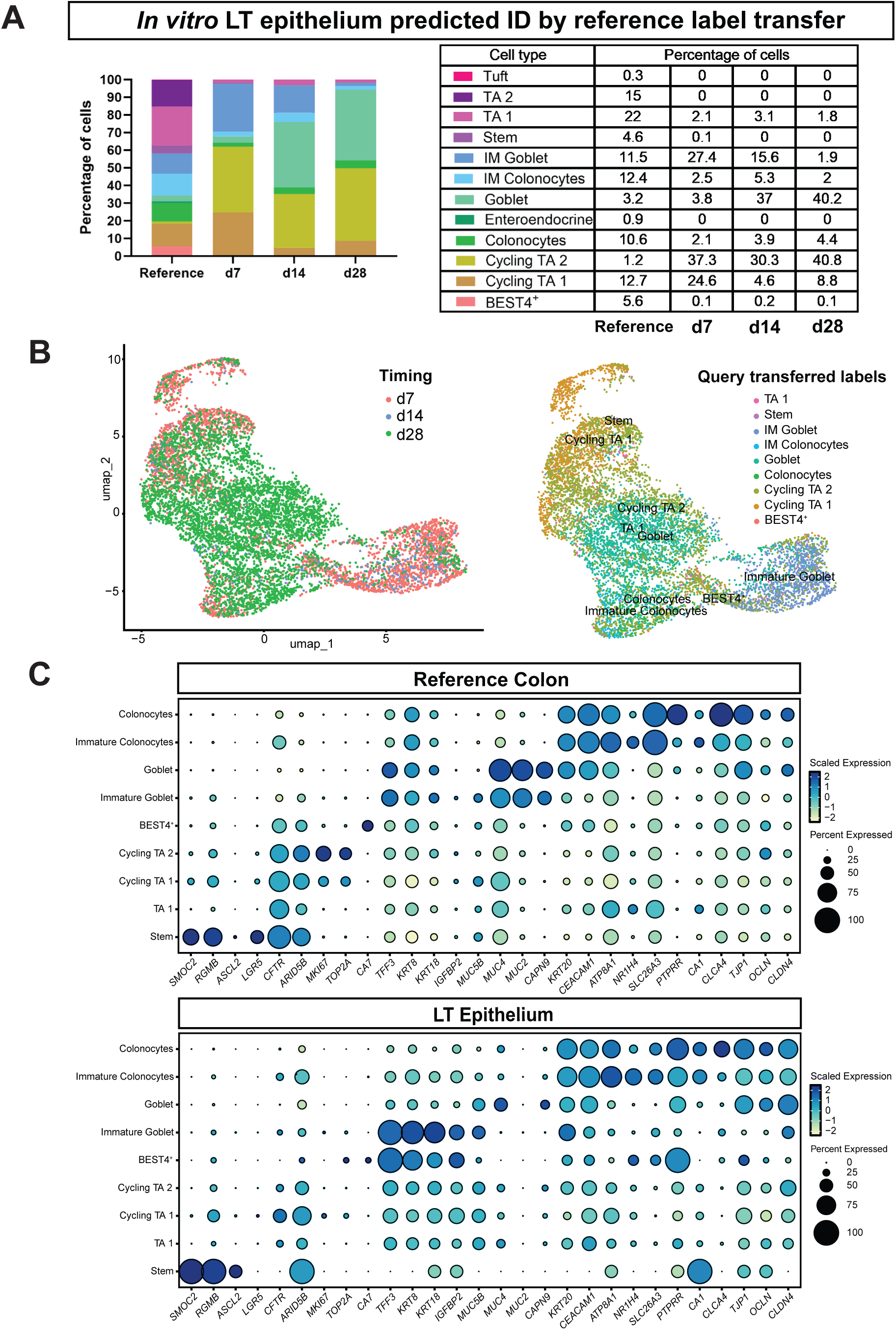
snRNA-seq of primary human colonoid derived LT epthelium across time with colon tissue reference label transfer identifies cell populations of the colonic epithelium *in vitro*. **A.** Cell type proportions of Colon 88 LT epithelium at day 7 (n = 3360 total cells), day 14 (n = 416 total cells) or day 28 (n = 4605 total cells) time points and reference ascending colon (n = 72,269 total cells). **B.** UMAP clustering of LT epithelium predicted ID by reference label transfer from colon tissue. UMAP of LT epithelium across time (left) and UMAP with query transferred labels (right). Immature (IM). **C.** Dot-plots representing the reference colon tissue (top) and LT epithelium (bottom) display gene marker expression levels of identified cell groupings.

### Continuous TEER measurements during LT epithelial barrier challenge reveal unique barrier responses for several days

We were interested to see how the LT epithelium model would respond to agents known to disrupt the epithelial barrier in a high-throughput format and if it was possible to study barrier dynamics with repeated exposure to these agents. We therefore established LT epithelial monolayers into 96 well transwell plates and started continuous TEER measurements using the electric cell-substrate impedance sensing (ECIS) TEERZ system beginning on day 2 or day 4 of LT monolayer culture (**Supplementary Figure 1C**). On day 8, TEER was measured and wells were distributed across experimental groups in order to balance the mean TEER across six control or treatment groups: 1. No treatment control, 2. basolateral TNF-α (5 ng/mL), 3. basolateral IFN-γ (5 ng/mL), 4. basolateral TNF-α (5 ng/mL) and IFN-γ (5 ng/mL) combined [56, 57], 5. apical TcdB (10 ng/mL) [58, 59], or 6. apical EDTA (2 mM; positive control for loss of barrier function) [60]. Continuous TEER measurements were paused while treatments were added, after which continuous TEER measurements resumed, applying fresh media plus appropriate treatment to the culture every other day for >1 week (**Figure 3A**). To examine the LT epithelium barrier responses to treatment, we calculated the percent change of mean daily TEER from the control group (**Figure 3B**). LT epithelium derived from three (n = 3) donor colonoid lines show unique donor-specific barrier function responses measured by significant changes in TEER (**Figure 3Bi, 3Bii, 3Biii**) during the treatment period while sustaining an intact cell layer (**Figure 3C**).

**Figure 3.**
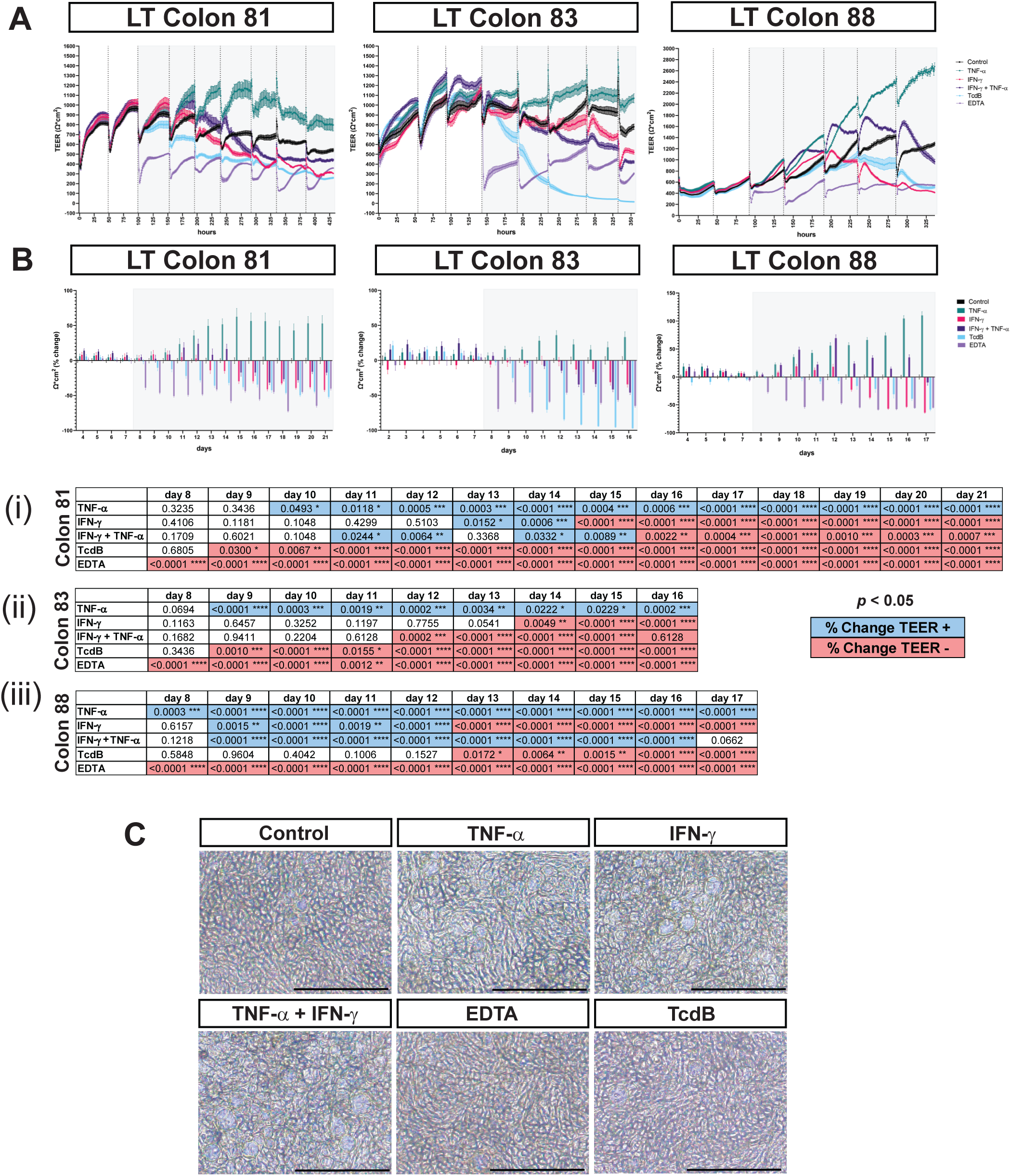
LT epthelium exposure to barrier challenge agents with continuous TEER measurement shows tolerant and intolerant barrier responses to repeated challenge over time. **A.** Hourly TEER graphed as mean ± SEM of each group with the following technical replicates: Control (n = 14), 5 ng/mL IFN-γ (n = 9), 5 ng/mL TNF-α (n = 9), 5 ng/mL IFN-γ and 5 ng/mL TNF-α (n = 9), 2 mM EDTA (n = 9), and 10 ng/mL TcdB (n=9). Treatments were applied to three different cell lines using Colon 81 (left), Colon 83 (middle), and Colon 88 (right). Treatment was added to respective groups at time indicated by dotted lines within gray shaded area of each graph. **B.** Daily mean percent change in TEER from control ± SEM of each group per cell line. Bi. Bii. Biii. Statistical significance of percent change of mean daily TEER from control was determined using *p* values calculated by Welch’s *t*-test. A significant increase (blue *p* value) or decrease (red *p* value) in the percent change in TEER from control is shown. **C.** Representative bright field images of Colon 81 LT monolayer treatment groups at day 12. Images were taken at 10X with an Olympus IX73 microscope. Scale bars represent 100 μm.

TNF-α treatments produced a significant daily increase in TEER starting early in the treatment period for all three donor-specific LT epithelium cultures, called ‘Colon 81’, ‘Colon 83’ and ‘Colon 88’ (**Figure 3Bi, 3Bii, 3Biii**). Colon 83 exhibited a less pronounced increase in TEER with TNF-α treatment compared to Colon 81 and Colon 88 (**Figure 3Bii**). Colon 88 had an immediate significant increase in TEER beginning the first day of TNF-α treatment and continued through the last day of TEER measurement (**Figure 3Biii**).

At timepoints early in the IFN-γ treatment period, all three-donor specific LT epithelium barriers tolerated this cytokine, initially showing no significant change in TEER (**Figure 3Bi, 3Bii**). Only after repeated addition of IFN-γ did we measure significantly increased barrier permeability (**Figure 3Bi, 3Bii, 3Biii**). For example, initially Colon 81 and Colon 83 with IFN-γ treatment show no significant change in TEER from control (**Figure 3Bi, 3Bii**). After three treatments of IFN-γ, Colon 81 had a two-day period with a significant increase in TEER followed by a significant decrease in TEER at remaining time points (**Figure 3Bi**). Colon 88 produced a significant increase in TEER early in the IFN-γ treatment period compared to the other two lines, but after repeated addition of IFN-γ we measured a significant decrease in TEER later in the treatment period (**Figure 3Biii**).

IFN-γ and TNF-α in combination has been shown in previous epithelial monolayer studies to quickly increase barrier permeability compared to IFN-γ alone [56, 61]. However, Colon 81 and Colon 83 treated with IFN-γ and TNF-α tolerated the combination treatment at early time points, similar to the response to IFN-γ alone (**Figure 3Bi, 3Bii**). After five days of TNF-α, IFN-γ, and the combination treatment we observed structural changes to the Colon 81 epithelium although there was either no significant change in TEER from the control (IFN-γ) or significant increase in TEER from the control (TNF-α, TNF-α + IFN-γ) (**Figure 3Bi, 3C**). Interestingly, Colon 88 maintained significantly increased TEER in the presence of the cytokine combination (**Figure 3Biii**). This result was somewhat surprising, although recent literature has shown that IFN-γ signaling can drive epithelial TNF-α receptor-2 (*TNFRSF1B*) expression during colonic tissue repair. That is, in specific contexts IFN-γ and TNF-α may work together in a pro-repair/pro-regenerative capacity [62].

The *Clostridium difficile* cytotoxin TcdB produced reductions in TEER 3 hours after the first addition to Colon 81 and 12 hours after the first addition to Colon 83 (**Figure 3A**). The percent change in mean daily TEER from the control was significantly decreased the day following the first TcdB addition (**Figure 3Bi, 3Bii**). However, at early time points Colon 88 tolerated TcdB, with no significant differences in TEER for the first five days of the treatment period, followed by significant decreases in TEER after three repeated additions of TcdB to the culture (**Figure 3Biii**). As expected, all three-donor LT epithelial barriers had a significantly reduced TEER from the start of EDTA treatment to the last day (**Figure 3Bi, 3Bii, 3Biii**). Although Colon 81 TEER was significantly reduced in TcdB and EDTA treatment groups, an intact epithelium remained on the transwell as shown in representative images five days into the treatment period (**Figure 3C**), indicating that the LT epithelium did not undergo cell death in the presence of these potent barrier disruptors.

### High-content image analysis of LT epithelium post-barrier challenge identifies quantifiable treatment-specific cellular features

To evaluate the phenotypic consequences of the LT epithelial perturbation, we used high-content imaging and analysis to compare the treatment groups. We fixed the Colon 83 LT epithelium and stained with DAPI (DNA), HCS CellMask Green (whole cell), Zonula Occludens-1 (ZO-1) antibody, and Claudin-2 (CLDN2) antibody (**Figure 4A**) at the end of our experiment using repeated treatments with TNF-α, IFN-γ, TNF-α and IFN-γ combined, TcdB, EDTA, or no treatment (Control). Each treatment induced a distinct cellular phenotype as observed by high-content imaging of the CellMask staining (**Figure 4B**). Combined treatment with TNF-α plus IFN-γ resulted in globally intensified CellMask signal and characteristic valleys of dim intensity between neighboring cells, suggestive of altered cell spreading (**Figure 4B**). TcdB led to prominent regions of cell loss, while EDTA caused scattered detachment of individual cells, indicating compromised adhesion (**Figure 4B**).

**Figure 4.**
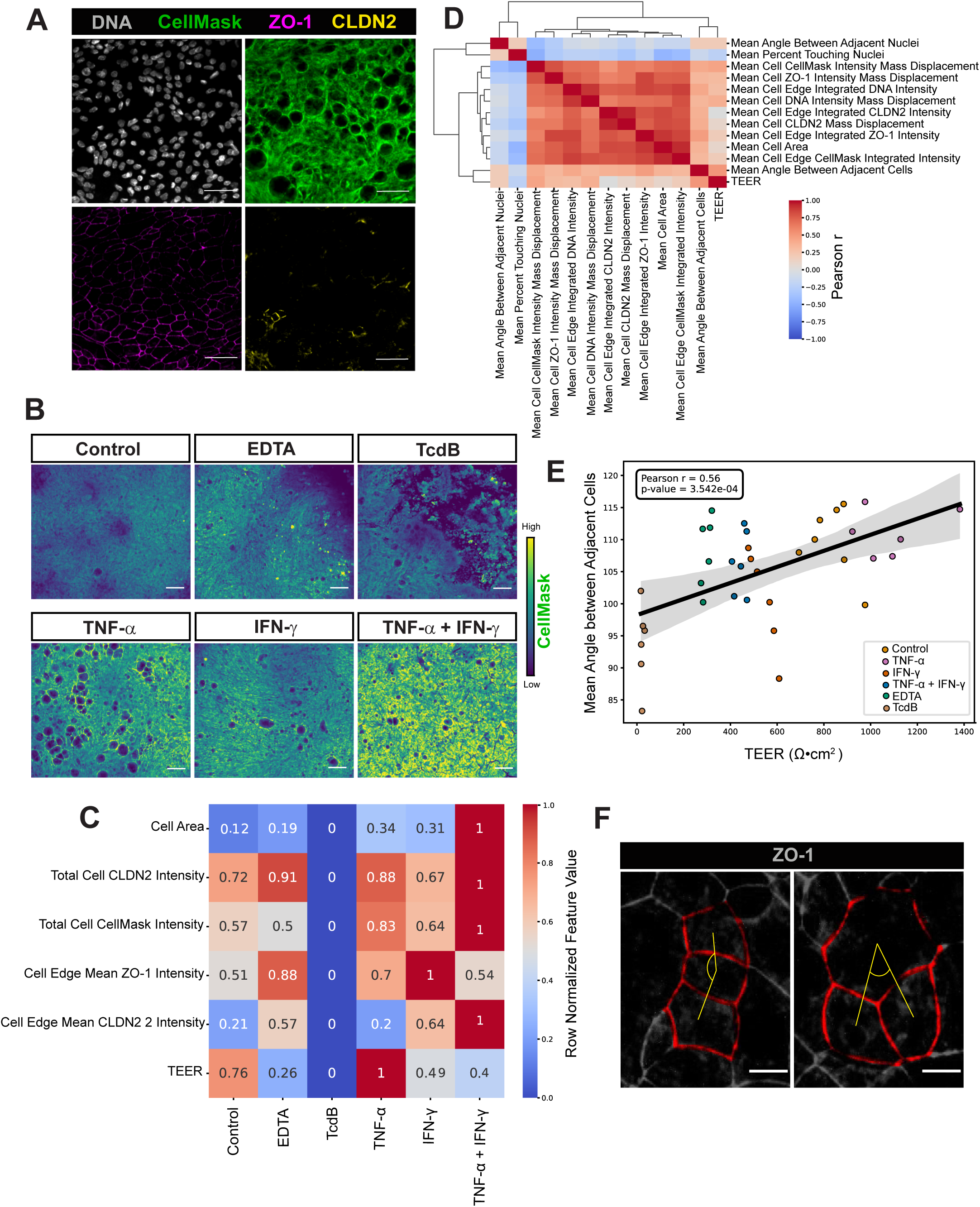
High-content image analysis of Colon 83 LT epithelium post-barrier challenge reveals cellu-lar features unique to treatment. All images were taken at 20X with a Yokogawa CellVoyager CQ1 Benchtop High-Content Analysis System. **A.** Representative high-content images of LT transwell epithelium IF staining DAPI (DNA), HCS CellMask Green, Zona Occludens-1 (ZO-1), and Claudin-2 (CLDN2). Scale bars represent 50 mm. **B.** Representative field-of-view images from treatment groups colored by CellMask Green intensity. Scale bars represent 100 mm. **C.** Normalized feature heatmap representing aggregated feature variation (row normalized feature value) between each treatment group (n=6 per group). A value of 1 indicates a positive relationship and 0 indicates no relationship. **D.** Clustermap of image-level feature correlations (n=16 per group). **E.** Top-correlating well-level aggregated imaging feature (most significant correlation with TEER): angle between adjacent cells against TEER. (n=6 per group). **F.** Schematic representation of how the angle is mea-sured between adjacent cells (red), where the center of each cell serves as the vertex and the two nearest adjacent cell centers define the vectors that form the angle (yellow). All scale bars represent 10 mm.

To quantify these differences, we visualized feature values across all treatments (**Figure 4C**). All cytokine-treated conditions, TNF-α, IFN-γ, TNF-α and IFN-γ combined, exhibited increased mean cell area, consistent with cytoskeletal rearrangement or hypertrophy. These treatments also led to progressively higher total CellMask and CLDN2 intensities, with the combination of TNF-α and IFN-γ combined eliciting the strongest response. CLDN2 edge intensity was highest with TNF-α and IFN-γ combined, moderately elevated in IFN-γ and EDTA, and unchanged in TNF-α relative to control. ZO-1 edge intensity increased in TNF-α and IFN-γ alone conditions but remained at control levels in the TNF-α and IFN-γ combined treatment group. TcdB treated wells consistently exhibited the lowest values for all image-derived features, consistent with significant cell loss and epithelial disruption. In contrast, EDTA treated barriers displayed a distinctive phenotype. Though TEER was reduced relative to control, morphological features such as cell area and total CellMask intensity were maintained. Notably, EDTA treatment induced elevated edge-localized ZO-1 and CLDN2 intensities and total CLDN2 intensity, indicating junctional reorganization rather than widespread detachment. Altogether, the analysis demonstrates that treatments produce specific and quantifiable phenotypes for different modalities of barrier disruption.

To assess the relationship between morphological features and barrier integrity, we calculated correlations between each image-derived metric and TEER measurements across all conditions. A clustermap of image-level feature correlations revealed several statistically significant positive correlations, with the mean angle between adjacent cells emerging as the most strongly associated feature (**Figure 4D**). This angular metric was a significant feature that clustered closely with TEER under all conditions (**Figure 4E, 4F**), with other significant features varying with TEER in treatment-dependent manner (**Supplementary Figure 3**). Overall, our high-content image analysis provided multiple quantitative outputs that identified cellular features unique to each treatment group and offered correlations of these features to barrier function via respective endpoint TEER measurements.

## Discussion

We present methods that enable a LT *in vitro* model of the primary human colonic epithelium on a transwell for 30 days, and we measured barrier function in real-time for up to 14 days. While several elegant approaches including microengineered collagen scaffolds [63] and organ-on-a-chip systems [64, 65] are available to interrogate the intestinal epithelium, the model developed here is constructed using simple, commercially available culture materials that do not require engineering expertise, and can be scaled for high-throughput assays with functional readouts including transporter function [66] or secretory responses [67]. We demonstrate quantitation of continuous TEER and high-content imaging of experimental endpoints using IF staining in a single-cell resolution and quantitative pipeline to assess epithelial barrier plasticity and integrity in a 96-well transwell plate. The formation of a confluent epithelium with functional barrier forms rapidly, within 72 hours of seeding (**Supplementary Figure 1C**). After formation of a stable barrier, we characterized LT epithelium histologically showing columnar cell morphology, proliferating cells, and a junctional barrier with quickly developing and sustained TEER that is more representative of the *in vivo* crypt compared to ST epithelial transwell method (**Figure 1**). Although terminally differentiated, ST epithelium average cell height was most similar to the *in vivo* crypt-base cell heights (**Figure 1B**) and measures similar sizes as immature infant-derived gastrointestinal epithelium on a transwell [68]. The LT epithelium cell heights overlap with each of the *in vivo* crypt regions; the crypt-base, the crypt-middle and crypt-top cells, better matching the morphology of the intact colonic crypt epithelial cell. Cell populations from crypt regions were identified by snRNA-seq with reference label transfer and indicated goblet and colonocyte lineages that mature over time, likely supported by TA-like populations of progenitor cells. (**Figure 2**) [69].

The LT system described here uses a transwell with media in both the apical and basal chambers; however, work from others has described an alternative transwell colonic epithelium with an air-liquid interface that establishes a LT monolayer [33, 34, 70]. While the nuanced difference between these two systems are unknown at this time, and would require a head-to-head comparison, it is important to point out that a liquid medium in the apical chamber is required for continuous TEER measurement that were conducted in this study. On the other hand, frequent medium changes in the apical chamber are required for the maintenance of the LT epithelium culture described here, and may disrupt the formation of a mucus layer, a key component of the colonic epithelial barrier that can be modeled *in vitro* [48, 71, 72]. Building a mucus layer in either LT system may be possible with the addition of VIP [10, 48]. Continuous TEER measurement of 96 well transwells containing early LT epithelium derived from three healthy donors subjected to barrier challenge with cytokines TNF-α, IFN-γ, TNF-α and IFN-γ combined initially show barrier responses to these cytokines in a tolerant way, without negative effects on TEER. This was unexpected given the observed structural changes mid-treatment period (**Figure 3C**), and the fact that these cytokines are generally assumed to be pro-inflammatory [73, 74]. Importantly, we observed donor-specific barrier responses that indicate donor-to-donor variation in the response to barrier-modifying agents. It is possible that barrier tightening recorded from the LT epithelium using Colon 81 and Colon 88 treated with TNF-α and IFN-γ combined may be responding in a pro-regenerative manner, given that IFN-γ functions as a pro-repair molecule by increasing TNF-α receptor 2 (*TNFRSF1B*) expression on intestinal epithelial cells following injury, and allowing TNF-α signaling to play a regenerative role *in vitro* and *in vivo* [62, 75].

Lastly, we performed IF staining followed by high-content image analysis of Colon 83 LT epithelium to determine if we could use high-content single-cell resolution phenotyping to interrogate the barrier integrity after repeated treatment. This analysis captures quantitative features of every cell in the culture dish, revealing that each treatment induces a distinct and measurable phenotype such that TcdB causes overt epithelial loss, EDTA disrupts junctional organization while preserving overall cell coverage, and cytokines promote coordinated increases in cell size and junctional protein abundance in a treatment-specific manner (**Figure 4**). Our image analysis captures the tight junction response to disrupting agents. CLDN2 is upregulated at the luminal crypt during inflammation and associated with increased permeability [54, 76]. We found a strong increase in CLDN2 intensity with the TNF-α and IFN-γ combined treatment (**Figure 4C**). Additionally, we detect an increase in cell area after all repeated cytokine treatments, with TNF-α and IFN-γ combined treatment having the greatest influence on cells size (**Figure 4C**). This may be an observation of tight junction remodeling that becomes leaky due to acto-myosin contractility [77]. While multiple features were correlated with TEER, the treatment-specific context of these correlations indicates that no single imaging feature reliably predicts TEER across all perturbations (**Figure 4E**, **Supplementary Figure 3**). Instead, TEER reflects a composite outcome of structural remodeling, where metrics such as cell area, junctional protein distribution, and cell alignment contribute in mechanistically distinct ways depending on the nature of the perturbation (**Figure 4C**).

Taken together, the data presented here outline new methods to establish a simple, scalable, and durable long-term human colonic barrier model that unites continuous TEER with single-cell, high-content phenotyping to resolve donor-specific and perturbation-specific responses over weeks, transforming the primary human intestinal epithelium into a practical discovery tool for mechanistic biology, disease modeling, and high-throughput therapeutic testing. This work lays the groundwork for precision, patient-matched studies and co-culture assays that connect epithelial structure to function at scale.

## Methods

All authors had access to the study data and had reviewed and approved the final manuscript.

### Primary Adult Human 3D Colonoid Culture

Primary colonoid cultures (**Table 1**) were established with isolated crypts from normal adult human ascending colon resections (University of Michigan Institutional Review Board HUM00105750, Not Regulated) [6, 69, 78]. Note, these lines were recently made available commercially (Millipore-Sigma Gastrointestinal Organoid Biobank). All established colonoid lines (University of Michigan Translational Tissue Modeling Laboratory Core Facility RRID:SCR_027333; <u>Biorepository</u> IRB: REP000000105) were submitted for a Human Comprehensive Clear Panel (Charles River Laboratories). Only Colon 88 tested positive for Human Herpes Virus 7. All other colonoid lines used for this study were virus panel negative. Additionally, all colonoid lines were authenticated as Short Tandem Repeat matches to their donor source tissue (STR; ATCC 135-XV) and confirmed colon with *SATB2* and *PDX1* qPCR analysis. Colonoids were cultured in six well plates (USA Scientific CC7682-7506) encapsulated in Matrigel (>9 mg/mL protein; <1.5 U/mL Endotoxin; Corning, 354254) diluted to 8 mg/mL with colonoids plus culture medium, plated in 25 x 10 μL droplets per well, and grown in a 5% CO_2_ and 37°C incubator. The culture medium was 10% IntestiCult Organoid Growth Medium (STEMCELL Technologies, 06010) with added Y-27632 (10 μM, Tocris Bioscience 1254) and Primocin (100 μg/mL, InvivoGen ant-pm-05) plus 90% Human Colonoid Medium [3, 6, 32, 78, 79] (HCM; 50% L-WRN conditioned medium (produced by L-cell line, ATCC CRL 3276) plus 50% 2X base medium—Advanced DMEM/F12 (Gibco, 12634028) with penicillin/streptomycin (25 U/mL/25 μg/mL, Gibco 15070063), GlutaMAX (4 mM, Gibco 35050061), HEPES (20 mM, Gibco 15630080), *N-*Acetyl-L-cysteine (2 mM, Sigma A9165), N-2 Supplement (2X, Gibco 17502048), and B-27 Supplement minus vitamin A (2X, Gibco 12587010)) with added recombinant human Epidermal Growth Factor (EGF; 100 ng/mL, R&D Systems 236-EG), Y-27632 (10 μM), A83-01 (500 nM, Tocris Bioscience 2939) SB 202190 (10 μM, Tocris Bioscience 1264), and Primocin (100 μg/mL). Culture medium was made weekly and replaced on the colonoid culture daily [80]. Colonoid wells were regularly tested negative for *Mycoplasma*, which included the removal of Primocin from the culture medium for 3 days and collecting medium from the colonoid well on day 3 for the assay (Lonza LT07318 and SouthernBiotech 1310001). Colonoids were passaged every 6-7 days by cold mechanical dissociation. The Matrigel droplets containing colonoids were scraped off the dish in 2 mL of cold Dulbecco’s Phosphate Buffered Saline (DPBS; Gibco 14190-144) with a Cell Lifter (Corning, 3008). The colonoid suspension was vigorously triturated 50 times with a Bovine Serum Albumin (BSA; 0.1% in DPBS, Sigma A8806) coated 1 mL pipette (Thomas Scientific 1159M42) in a BSA coated conical (5 mL, Dot Scientific T2540) and centrifuged at 300 x*g* for 3 minutes at 4°C. Pelleted colonoid fragments were triturated 30 times with a BSA coated 200 µL pipette in cold culture medium with CHIR99021 (2.5 µM, Cayman Chemical 13122), transferred into cold Matrigel to 8 mg/mL, resuspended 15 times and then plated in 10 μL droplets. The culture dish with colonoid droplets was placed in a 5% CO_2_ and 37°C incubator for 10 minutes to allow Matrigel to solidify, followed by adding 2 mL of 10% IntestiCult and 90% HCM plus CHIR99021 (2.5 µM) pre-warmed in a 37°C water bath to the droplet wells. DPBS plus antibiotics, Primocin (100 μg/mL) and Gentamicin (10 μg/mL, Gibco 15710064), was added to empty wells to humidify the plate. CHIR99021 was removed from the culture medium the day after passaging. Medium volumes in colonoid wells were increased post-passage to support new growth over time (2 mL day 1-2, 2.5 mL day 2-4, 4 mL day 4-6 or 7).

**Table 1.**

| **Normal Colonoid Lines and Source Tissues** |  |  |  |  |
| --- | --- | --- | --- | --- |
| **ID** | **Location** | **Age** | **Sex** | **Source** |
| Colon 81 | Ascending Colon | 49 | F | TTML-University of Michigan |
| Colon 83 | Ascending Colon | 45 | F | TTML-University of Michigan |
| Colon 88 | Ascending Colon | 33 | F | TTML-University of Michigan |
| Colon 87 | Ascending Colon | 21 | M | TTML-University of Michigan |
| Colon 89 | Ascending Colon | 55 | M | TTML-University of Michigan |

TTML: Translational Tissue Modeling Laboratory Biorepository IRB: REP000000105, all specimens HUM00105750 Not Regulated

### Seeding dissociated 3D colonoids onto 2D transwell membranes

Colonoids were expanded from low passage cryopreserved stocks up to passages 9-30. 6-7 days post passage, colonoids had mostly cystic morphology and high density (**Supplementary Figure 1A**). Approximately 6 hours before colonoid dissociation (day - 1), the culture medium was refreshed. Colonoids were dissociated into small cell aggregates using a modified protocol from previously published methods [22, 25]. 1 well of a 6-well dish containing 25 x 10 µL Matrigel droplets of colonoids were collected in 2 mL of EDTA (0.5 mM in DPBS, Invitrogen 15575020) using a Cell Lifter. Next, colonoids were gently triturated with a 1 mL pipette five times against the bottom of a conical (15 mL, Fisher Scientific 14-955-237) brought up to 10 mL in EDTA, and centrifuged at 200 x*g* for 5 minutes at 4°C. The colonoid pellet was gently triturated with a 1 mL pipette five times in 1 mL of Trypsin-EDTA (0.05%, Gibco 25300054) and Y-27632 (10 μM) at room temperature. Immediately, the conical was capped and moved into the 5% CO_2_ and 37°C incubator for 3.75 minutes. Quickly, 1 mL of deactivating serum containing Advanced DMEM/F12, Fetal Bovine Serum (10%, Corning 35-015-CV), HEPES (10 mM) and Y-27632 (10 μM) was added to trypsinized colonoids and triturated vigorously with a 1 mL pipette 50 times at room temperature. Next, colonoid aggregates were passed into a 50 mL conical (Corning 430290) through a cell strainer (40 µM, Corning 352340) pre-wet with 1 mL of deactivating serum and then rinsed with 1 mL of deactivating serum for every 1 mL volume of aggregates strained. Aggregates ≤40 µM were spun at 400 x*g* for 5 minutes at 4°C. The pellet was gently resuspended in 1.05 mL HCM plus CHIR99021 (2.5 µM) seven times with a 1 mL pipette at room temperature. Next, the 50 mL conical of aggregates was swirled and 50 μL of the suspension was collected for counting using a Multisizer 4e Coulter Counter (Beckman Coulter), set to count 6-60 µm sized particles. Counted aggregates were diluted further in HCM plus CHIR99021 to 200,000 per 100 µL and seeded on the apical side of Collagen IV (CIV; 1 mg/mL in 100 mM Acetic acid (Fisher Scientific A38-212) to 33.3 µg/mL in UltraPure distilled water (Invitrogen 10977-015), Sigma-Aldrich C5533) coated PET transwell membranes (24 well, Costar 3740 or 96 well, STEMCELL Technologies 100-0419) that were warmed in the 37°C incubator for 3 hours prior to seeding. The excess CIV was aspirated from the apical side of the membrane, colonoid aggregates in suspension were swirled, and 100 µL of suspension was pipetted per 0.33 cm^2^ membrane or 43.3 µL pipetted per 0.143 cm^2^ membrane. Room temperature HCM plus CHIR99021 was added to the basolateral side of the membrane or directly into the well of the culture dish (600 µL per 24 well or 235 µL per 96 well) (**Supplementary Figure 1B**). “Edge effect,” or the evaporation of medium from outer most wells of the culture dish was strong within transwell culture dishes. For all experiments, transwells were not cultured in wells on the outer edge of any culture plate. These wells were filled with DPBS plus antibiotics. Lastly, seeded tranwells with DPBS humidified outer wells were carefully transferred to the 5% CO_2_ and 37°C incubator and left undisturbed for 24 hours.

### Long-term (LT) epithelium 2D transwell culture

24 hours post seeding (day 0), the medium in the on the apical (200 µL per 24 well or 75 µL per 96 well) and basolateral (600 µL per 24 well or 235 µL per 96 well) sides of the transwell were changed to HCM, omitting SB 202190 and CHIR99021. 48 hours post seeding (day 1), the medium in the apical compartment was changed to 2D Differentiation Medium [24, 25] (2DM; 50% 2X base medium plus 50% Advanced DMEM/F12 with recombinant human EGF (50 ng/mL), recombinant human NOGGIN (50 ng/mL, R&D Systems 6057-NG), human Gastrin I (10 nM, Sigma-Aldrich G9020), and Primocin (50 μg/mL)) with Y-27632 (2.5 μM). 72 hours post seeding (day 2), the media in the apical side was changed to 2DM without Y-27632 and the HCM without SB 202190 on the basolateral side was refreshed. These apical and basolateral media conditions were maintained throughout long-term culture and changed every other day. A summarized schematic of the protocol and representative images of cell cultures during the set up are provided in **Supplementary Figure 1C**.

### LT epithelium 2D transwell culture with barrier challenge

After recording a steady TEER measurement, this indicated that a functional epithelial barrier was established, and the barrier challenge began. The addition of cytokines for barrier challenge occurred on day 8 and included recombinant human TNF-α (5 ng/mL, InvivoGen rcyc-htnfa) and/or recombinant human IFN-γ (5 ng/mL, InvivoGen rcyec-hifng) diluted in the basolateral medium where the *Clostridium difficile* Toxin B (TcdB; 10 ng/mL, Sigma-Aldrich SML1153) and EDTA (2mM) were diluted in the apical medium. Basolateral or apical application was chosen to reflect where the epithelium would encounter the cytokines, TcdB, or EDTA *in vivo* and were added to transwell cultures with media changes. Bright field images were captured regularly using an Olympus IX73 microscope using the 10X objective.

### Transepithelial Electrical Resistance (TEER) measurements

TEER measurements started on LT epithelium culture day 1 or 2. The epithelial volt/ohm meter and electrode (World Precision Instruments EVOM2 and STX2) were used for manual TEER measurements on 24 well transwells. In the biological safety cabinet, the electrode was sprayed with 70% ethanol, dried, rinsed in warm DPBS, and gently shaken to wick away excess liquid. The transwell plate was placed on top of a warm pack (Fisher Scientific 03-531-53) pre-heated in the 37 °C incubator. Next, the electrode was held vertically and the long end was lowered through an opening in the transwell insert into the bottom of the plate well or the basolateral side of the epithelium, while the shorter end of the electrode remained on the apical side above the epithelium on the transwell membrane. The electrode was held in place for a few seconds to record the ohm value and then ohms were recorded from each remaining transwell in succession. Ohm values were recorded one time per day and always before the media on the well was changed. The TEER value was calculated for each transwell: ohm (Ω) x transwell surface area (0.33 cm^2^) = TEER (Ω*cm^2^). The ECIS TEERZ system (Applied BioPhysics) [81–83] was used to record and compute continuous TEER measurements from 96 well transwell plates inside the 5% CO_2_ and 37°C incubator. The measurement was briefly paused, transwell plate removed for media change, and then resumed one time every other day.

### Histology and Image analysis

Transwell epithelium was fixed with 10% neutral buffered formalin (Fisher Scientific 23-245685) for 15 minutes at room temperature, washed two times with DPBS, and the membrane with attached epithelium was removed from the transwell insert using a no. 11 scalpel blade (Aspen Surgical Products 371111). The circular epithelium with membrane was embedded in HistoGel (Epredia HG4000012) and placed in 70% ethanol in Milli-Q filtered water (100% Ethanol; Decon Labs 2701). Colon tissue was fixed with 10% neutral buffered formalin overnight, washed the next day, and placed directly in 70% ethanol. Paraffin processing, embedding, sectioning, and Hematoxylin & Eosin staining were performed at the University of Michigan Rogel Cancer Center Tissue and Molecular Pathology Shared Resource and University of Michigan Integrative Musculoskeletal Health Core Center Orthopaedic Research Laboratories. 40X brightfield digital representations of the glass microscope tissue sections were made with a Leica Aperio AT2 or Leica Aperio GT450 (Leica Biosystems Imaging, Inc.) at the Michigan Medicine Department of Pathology Digital Pathology Core. Images and cell heights were taken directly from the digital scan using Aperio ImageScope software v12.4.6.5003 (Leica Biosystems Imaging, Inc.). Cell heights were measured directly on the digital scan using the Ruler Tool and recorded into the Annotations box.

### Whole-mount Immunofluorescence (IF)

LT epithelium was kept attached to the insert and washed two times with warmed DPBS on the apical (200 µL per 24 well or 100 µL per 96 well) and basolateral (400 µL per 24 well or 200 µL per 96 well) sides. Next, the epithelium was fixed with cold methanol (Fisher Scientific A412-1) for 15 minutes at −20°C, washed three times with cold DPBS at room temperature for 5 minutes, and stored at 4°C. To begin immunolabeling, the fixed epithelium was incubated in blocking buffer composed of 3% BSA and 0.3% Triton X-100 (Sigma-Aldrich T9284) in DPBS for 1 hour at room temperature. Blocking buffer was removed and primary antibodies diluted in antibody buffer composed of 1% BSA and 0.3% Triton X-100 were added to the apical (100 µL per 24 well or 20 µL per 96 well) side with DPBS on the basolateral side. Antibodies were incubated on the epithelium overnight at 4°C. The next day, the epithelium was washed three times with DPBS for 15 minutes at room temperature. A secondary antibody with fluorophore was sometimes used when the primary antibody was not available with conjugated fluorophore. In this case, a secondary antibody was diluted in antibody buffer and incubated on the epithelium for 1 hour at room temperature. All antibodies and dilutions used can be found in **Table 2**. For 96-well transwells, HCS CellMask Green Stain (1:10,000, Invitrogen H32714) was added to the secondary antibody buffer. The epithelium was washed three times with DPBS for 5 minutes at room temperature. Lastly, nuclei were stained with DAPI in DPBS (1:10,000 of 10 mM stock in UltraPure water, Invitrogen D3571) for 5 minutes at room temperature and then washed three times with DPBS for 5 minutes at room temperature. Immunolabeling incubations and washes were performed on a slow rocking platform. For immunolabeled epithelium on transwell membranes in 24 well plates, the DPBS was removed from the apical side and using a no. 11 scalpel blade, the transwell membrane with attached immunolabeled epithelium was cut out of the transwell insert from the basolateral side. Microforceps (Fisher Scientific 08-953E) were used to handle the edge of the membrane and placed cell side up on a 50 µL droplet of ProLong Gold Antifade Mountant (Invitrogen P36930) in the center of a microscope slide (Alkali Scientific Inc. SM2575). Another 50 µL droplet of mountant was gently added to the top of the cells followed by a glass coverslip (Fisher Scientific 12-541B). Slides were dried overnight at room temperature, edges sealed with clear nail polish, and slides were stored at 4°C until imaging. Immunolabeled LT epithelium in 96 well plates were stored in DPBS at 4°C until imaging.

**Table 2.**

| **Whole-mount Immunofluorescence Antibodies** |  |  |  |
| --- | --- | --- | --- |
| **Antibody** | **Dilution** | **Source** | **Catalog #** |
| Occludin (OC-3F10) mouse monoclonal, AlexaFluor 488 | 1:200 | ThermoFisher Scientific | 339194 |
| Ki-67 (D3B5) rabbit monoclonal, AlexaFluor 647 | 1:100 | Cell Signaling Technology | 12075 |
| Mucin 2 (F-2) mouse monoclonal, AlexaFluor 594 | 1:50 | Santa Cruz Biotechnology | sc-515032 |
| E-cadherin (36/E-Cadherin) mouse monoclonal, AlexaFluor 488 | 1:50 | BD Biosciences | 560061 |
| ZO1 (ZO1-1A12) mouse monoclonal, AlexaFluor 594 | 1:200 | Cell Signaling Technology | 12075 |
| Claudin-2 (MH44) rabbit polyclonal | 1:100 | ThermoFisher Scientific | 339194 |
| Alexa Fluor 647 AffiniPure Donkey Anti-rabbit IgG (H+L) | 1:500 | ThermoFisher Scientific | 51-6100 |
| Ki-67 (D3B5) rabbit monoclonal, AlexaFluor 647 | 1:100 | Jackson ImmunoResearch | 711-605-152 |

### Confocal microscopy and image processing

Slides were imaged with a Nikon A1 light scanning confocal microscope using a 40X objective with oil immersion. Maximum intensity projections acquired from 0.24 µm Z-stack images and orthogonal view images were created using Fiji (National Institutes of Health, USA) [84].

### High-Content Imaging, Processing, and Cell Segmentation

96-well transwell plates were imaged on the Yokogawa CellVoyager CQ1 Benchtop High-Content Analysis System using a UPLXAPO 20x/.8 NA dry objective, and maximum intensity projections were acquired from 1.3µm Z-stack images. Cell nuclei were segmented using Cellpose 3.0 from DAPI images and individual cells were segmented using a custom Cellpose 3.0 model trained on ZO-1 images [85, 86]. Image, cell, and nuclear measurements were acquired using CellProfiler 4.2.5 [87].

### Single nuclei isolation from LT epithelium and Single nucleus RNA sequencing (snRNA-seq)

LT-epithelium cultures were collected on days 7, 14 and 28 from the same set of transwells using Colon-88 at passage 9. Cells were gently scraped off the transwell membrane with a mini cell scraper (United Biosystems MCS-200) in 200 μL of cold DPBS and transferred to a cryovial (Fisher Scientific 12-567-501) containing 1.3 mL of cold DPBS with a cut 0.1% BSA coated 200 µL pipette tip and centrifuged at 500 x *g* for 3 minutes at 4°C. Excess DPBS was gently removed from the cell pellet and the pellet was immediately flash frozen in liquid nitrogen. Two transwell cell culture samples were collected per time point and pooled into the same cryovial for nuclei isolation. All samples were stored in liquid nitrogen until nuclei isolation. Nuclei were isolated using the Chromium Nuclei Isolation kit (10x Genomics 1000493) and protocol for Single Cell Gene Expression and Chromium Fixed RNA Profiling (10x Genomics CG000505 Rev A) with added RNase inhibitor (2 U/mL, Roche 3335402001) throughout the isolation. Because this protocol is designed for isolation of nuclei from whole tissue, we modified the initial lysis steps for gentler dissociation of the cell pellets by slowly triturating the frozen cell pellet in cold lysis buffer 15 times with a 1 mL pipette and omitting the use of a pellet pestle. This was followed by a 5 minute incubation in the lysis buffer on wet ice with additional gentle trituration at 2.5 minutes and immediately after the 5 minute incubation. Isolated nuclei were resuspended in UltraPure BSA (10 mg/mL, Invitrogen AM2616) diluted in DPBS with RNase inhibitor, counted, and loaded into the 10x Chromium to create single-nuclei droplets targeting 5,000 nuclei. Single-cell libraries were prepared using the Next GEM Single Cell 3’ Gel Bead kit v3.1 (10x genomics PN-1000147) according to the manufacturer’s instructions. Illumina Novaseq performed all scRNA-seq. Library prep and next-generation sequencing was carried out in the Advanced Genomics Core at the University of Michigan.

### snRNA-seq data processing and analysis

Reads were mapped to the human genome (GRCh38-2020-A) and gene expression matrices were generated using the CellRanger (v7. 0) under standard parameters as determined by the University of Michigan Advanced Genomics Sequencing Core. Intact nuclei were re-determined on raw matrices using CellBender (v0.1) [88], which also corrected for ambient RNA at a false-positive rate of 0.01. CellBender corrected gene expression matrices were important into Seurat v4 [89] in RStudio v1.4 running R v4.4 for further analysis. This generated 11,735 nuclei for additional QC and analysis. All commands were run with default parameters unless otherwise noted.

#### Preprocessing/QC Filtering

Nuclei were removed from analysis if they did not meet the following QC thresholds: between 500 to 7000 features and less than 5% mitochondria RNA reads. This removed 3,354 low quality nuclei, leaving 8,381 high-quality nuclei for further analysis. For each individual count matrix values were log-normalized using Seurat::NormalizeData() and variable features were identified using Seurat::FindVariableFeatures().

#### Data Scaling and Principal Component Analysis

Common variable features amongst all datasets were determined by Seurat::SelectIntegrationFeatures(). Features from Seurat::SelectIntegrationFeatures() were then scaled using Seurat::ScaleData() and principal components determined by Seurat::RunPCA().

#### Data Integration, Dimensional Reduction, Clustering

Data was integrated using reciprocal PCA integration. Anchors were first determined using Seurat::FindIntegrationAnchors(). A batch-corrected assay (‘integrated) was generated using Seurat:IntegrateData() with 30 dimensions. The ‘integrated’ assay was scaled by Seurat::ScaleData() and Principal Components for the ‘integrated’ assay were determined by Seurat::RunPCA(). PCs 1-12 were used to generate a Uniform Manifold Approximation and Projection (UMAP) by Seurat::RunUMAP() and for nearest-neighbor graph construction by Seurat::FindNeighbors(). Louvain clustering was performed at a resolution of 0.4 using Seurat::FindClusters().

#### Inference of In Vitro Epithelial Cell Type Identities by Label Transfer

Normal adult colon tissue reference snRNA-seq data from https://datadryad.org/dataset/doi:10.5061/dryad.8pk0p2ns8 [39, 40] was loaded into R and a new UMAP was generated using PCs 1:30 calculated by the authors using with return model = TRUE. A set of anchors for label transfer were calculated between the colon reference to the *in vitro* 2D monolayer query using Seurat::FindTransferAnchors () on the ‘decontXcounts’ assay in the reference and the ‘RNA’ assay in the query. Labels were then transferred by Seurat::MapQuery().

#### Gene Expression Analysis

All gene expression analysis was conducted on the CellBender corrected log-normalized gene expression matrices stored under the ‘RNA’ assay. Significant differences in gene expression between clusters was determined using Seurat::FindMarkers() with cutoffs of log_2_ fold-change = 0.25 and detected in at least 25% of nuclei in one of the clusters being compared.

### snRNA-seq data and code availability

Sequencing data is available at EMBL-EBI ArrayExpress under accession number E-MATB-15263. The code used for the snRNA-seq analysis is available at https://github.com/jason-spence-lab/Cuttitta_et_al_2025.

### Statistical Analyses

The following statistical analyses were conducted using GraphPad Prism 10 (GraphPad Software, San Diego, CA). The significance of mean cell heights between sample groups (**Figure 1B**) and the significance of the percent change in mean daily TEER between treatment groups and the control group (**Figure 3B, Bi, 3Bii, 3Biii**) was calculated using Welch’s T-Test. Pearson correlation *p* values were calculated using scipy.stats version 1.10.1 (**Figure 4**, **Supplementary Figure 3**). Differences were considered statistically significant at *p* < 0.05.

## Abbreviations

2D: 2-dimensional;
2DM: 2D differentiation medium;
3D: 3-dimensional;
CLDN2: Claudin-2;
ECAD: E-cadherin;
ECIS: electric cell-substrate impedance sensing;
EDTA: ethylenediaminetetraacetic acid;
EGF: epidermal growth factor;
EVOM: epithelial volt/ohm meter;
IHC: immunohistochemical;
IF: immunofluorescence;
IFN-γ: Interferon gamma;
ISC: intestinal stem cell;
LT: long-term;
MKI67: Marker of proliferation Ki-67;
MUC2: Mucin 2; OCLN, Occludin;
ST: short-term;
TA 1: transient amplifying 1;
TA 2: transient amplifying 2;
TcdB: *Clostridium difficile* Toxin B;
TEER: transepithelial electrical resistance;
TNF-α: Tumor Necrosis Factor alpha;
VIP: Vasoactive intestinal peptide;
ZO-1: Zonula occludens-1

