## Supplementary Figures for "A scalable long-term *in vitro* model of human colonic epithelium with continuous barrier function"

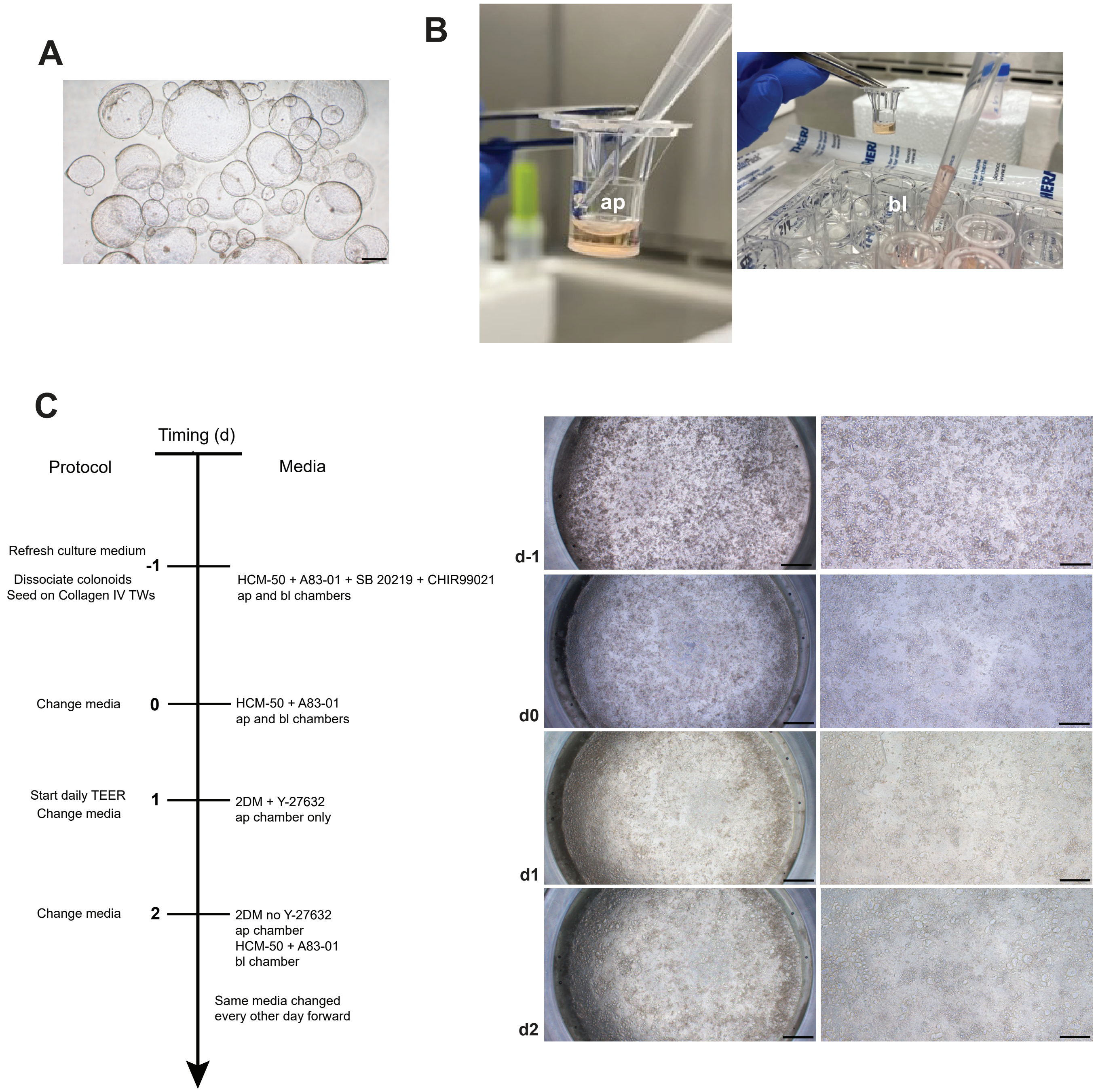


**Supplementary Figure 1. LT epthelium model components and protocol summary.**
**A**. Representative image of 3D colonoid culture at the time of harvest for LT epithelilum culture set up.
Shown is Colon 88 passage 14. Image was taken at 4X with an Olympus IX73 microscope.The scale bar represents 500 $\mu$m.
**B**. Apical (ap) and basolateral (bl) transwell compartments.
**C**. Schematic of protocol summary (left) with corresponding representative images of epithelial monolayer development (right).
Timing is noted by day (d). Images were taken at 4X (left image panel) and 10X (right image panel) with an Olympus IX73 microscope.The scale bars represent 500 $\mu$m (left image panel) and 200 $\mu$m (right image panel).

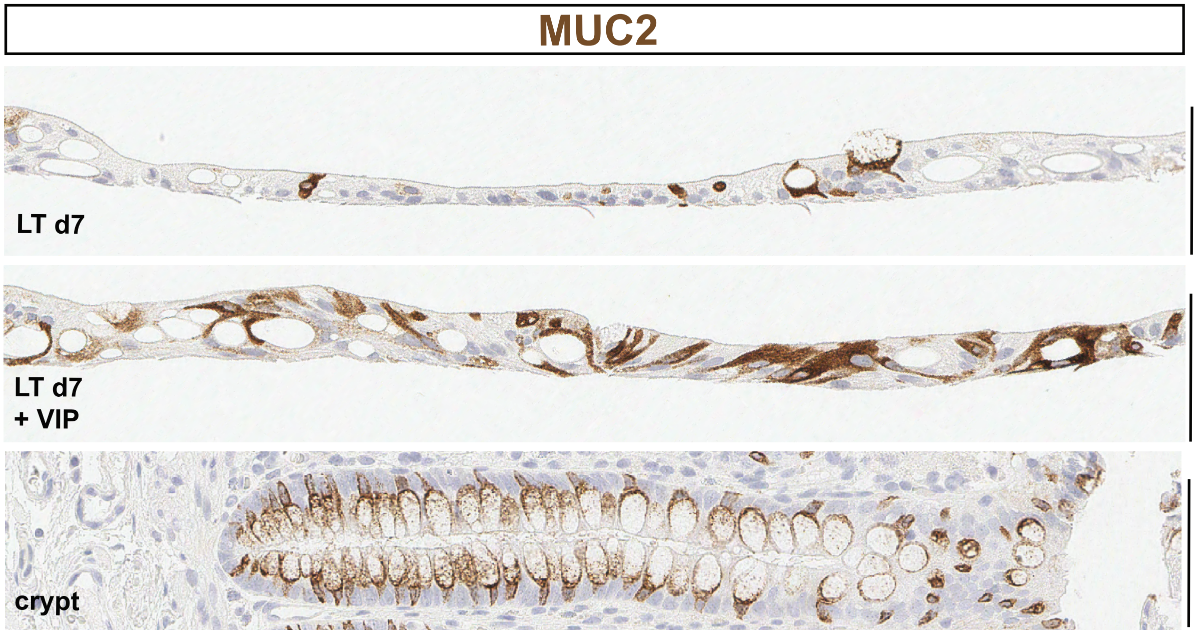


**Supplementary Figure 2. LT epthelium produces MUC2+ goblet cells.**
Representative images of histological cross-sections from *in vitro* LT epithelium fixed on day 7 (d7) and adult *in vivo* colon crypt with MUC2-DAB IHC labeled goblet cells. LT epithleium (top), LT epithleium + 330 ng/mL Vasoactive Intestinal Peptide (VIP) in basolateral culture medium starting day 1 (middle), and adult *in vivo* colon crypt (bottom). Images were taken at 40X with an Leica Aperio slide scanner. All scale bars represent 80 μm. IHC was performed at the University of Michigan Rogel Cancer Center Tissue and Molecular Pathology Shared Resource using mouse monoclonal MUC2 (MRQ-18) antibody (Sigma-Aldrich 291M-16) and a 1:500 dilution.

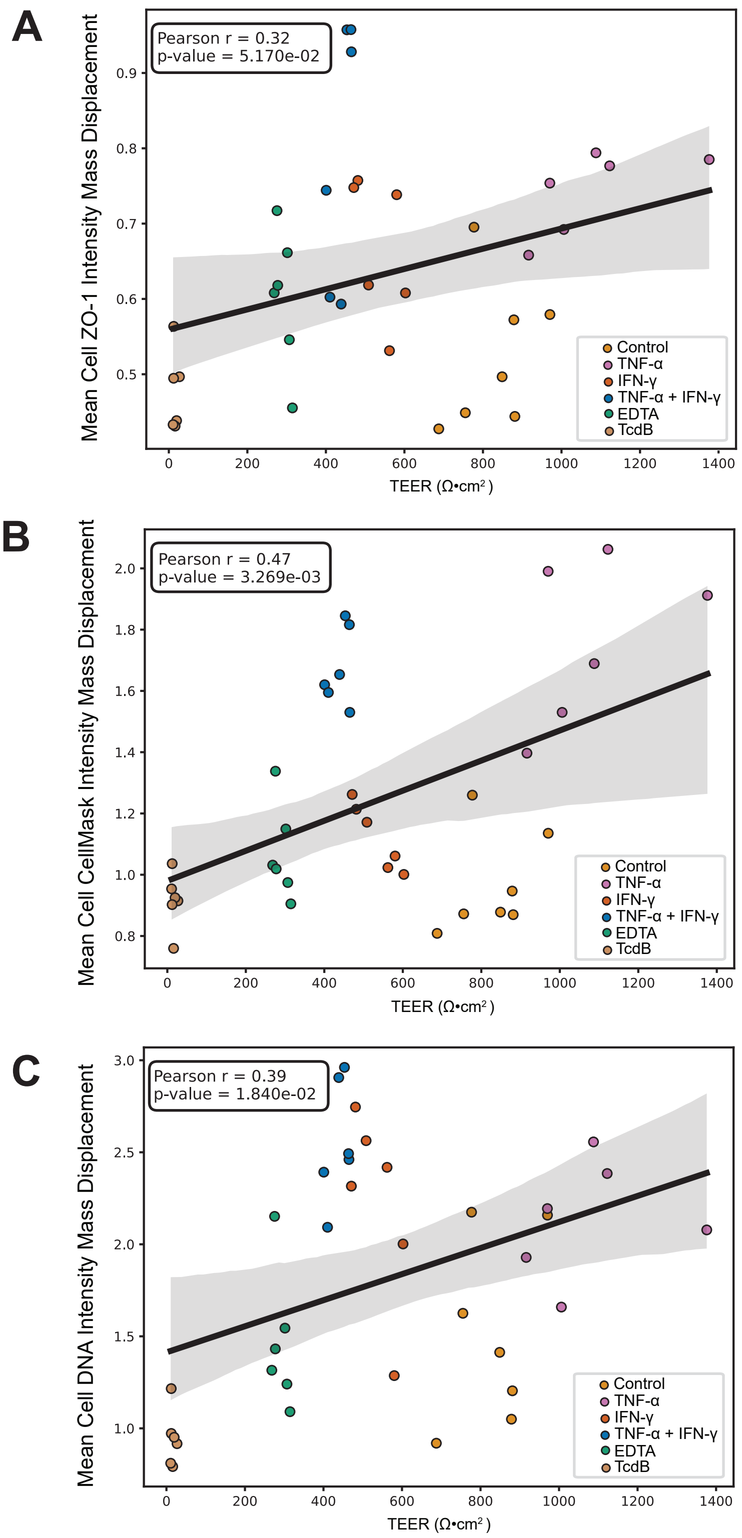


**Supplementary Figure 3. Additional top well-level aggregated imaging features from Colon 83 LT epithelium with significant correlation to TEER post-barrier challenge shows treatment dependent patterns.**

**A**. Mean cell ZO-1 intensity mass displacement (n=6 per group) clustering with TEER.

**B**. Mean cell CellMask intensity mass displacement (n=6 per group) clustering with TEER.

**C**. Mean cell DNA intensity mass displacement (n=6 per group) clustering with TEER.
